# Delineating the Role of ITGAM in Macrophage Dynamics and Cardiac Modulation during Sepsis-Induced Cardiomyopathy

**DOI:** 10.1101/2024.03.08.583788

**Authors:** Qinxue Wang, Haobin Huang

## Abstract

**Background:** Sepsis-induced cardiomyopathy (SIC) represents a critical complication of sepsis, characterized by reversible myocardial dysfunction and alterations. Despite extensive research, the molecular mechanisms underlying SIC remain poorly understood.

**Methods:** Utilizing bioinformatics analysis of RNA-seq and scRNA-seq data from the GEO database, we identified key immune cell populations and molecular markers associated with SIC. Our in vitro and in vivo studies focused on the roles of ITGAM and ICAM-1 in macrophage recruitment and transformation as well as the impact of these changes on cardiac function.

**Results:** Bioinformatics analysis revealed significant alterations in gene expression and immune cell composition in cardiac tissue during SIC, with macrophages being the predominant immune cell type. ITGAM was identified as crucial molecule in this process. In vitro experiments demonstrated an upregulation of ITGAM in macrophages and ICAM-1 in endothelial cells following LPS stimulation, indicating their roles in immune cell recruitment and interaction. Furthermore, the use of ITGAM-neutralizing antibodies led to reduced macrophages infiltration and initially improved cardiac function in SIC mice, but resulted in increased mortality in later stages. These findings highlight the dual role of ITGAM in SIC, facilitating early-stage inflammation and later-stage cardiac recovery.

**Conclusion:** This study elucidates the complex dynamics of immune cells in SIC, with a particular emphasis on the role of ITGAM in macrophage modulation. The findings provide new insights into the reversible nature of myocardial dysfunction in SIC and underscore the importance of targeted therapeutic strategies for effective sepsis management.

**Highlights:** Identifies ITGAM as a key modulator in macrophage dynamics during sepsis-induced cardiomyopathy (SIC).

Elucidates the impact of ITGAM on cardiac function in SIC.

Reveals new insights into the immune-cellular mechanisms in SIC pathology.

## Introduction

Sepsis, a life-threatening condition triggered by the body’s excessive and dysregulated response to infection, remains a global health challenge with alarmingly high mortality rates[1]. Among the myriad of complications associated with sepsis, sepsis-induced cardiomyopathy (SIC) has emerged as a significant concern[2]. Characterized by reversible myocardial dysfunction and alterations, SIC has garnered considerable attention in both scientific and clinical realms. Nonetheless, deciphering the intricate mechanisms underlying the injury-self-rehabilitation phenomenon still remain challenging[2].

The multifaceted nature of SIC has prompted in-depth studies into its pathophysiology, diagnosis, and management. While definitive evidence remains elusive, abnormalities in global or microcirculatory coronary function are posited as potential causative factors for SIC in some preliminary studies[3, 4]. Furthermore, inflammation, through its intricate disturbance involving Toll-like receptor signaling, cytokine production, and inducible nitric oxide synthase activation, precipitates alterations in cardiomyocyte contractility, oxidative stress, and myocardial dysfunction in SIC[5, 6]. Additionally, mitochondrial dysfunction, stemming from various mechanisms including mitochondrial swelling and mitochondrial DNA damage, plays a crucial role in SIC by disrupting energy production, leading to oxidative stress and potential structural damage[7, 8]. While prior studies have elucidated a comprehensive understanding of the molecular mechanisms underpinning SIC, given that SIC is inherently secondary to sepsis, further exploration into the roles of immune cells and pathways modulating cardiac function during sepsis warrants increased attention.

The emerging frontier of genetic lineage tracing and high-dimensional sequencing techniques have provided new opportunities for understanding the roles of different immune cell populations in the pathogenesis of SIC. In this research, we employed bioinformatics analysis to track immune cell populations and associated molecular markers in SIC. Subsequently, we substantiated these findings through cell and animal model validations. With a more profound understanding of the mechanisms underlying SIC pathogenesis, it is anticipated that we can develop increasingly effective, patient-specific therapeutic strategies, augmenting survival prospects while concurrently minimizing adverse effects.

## Materials and methods

### Data collection for bioinformatics analysis

RNA sequencing (RNA-seq) and single-cell RNA sequencing (scRNA-seq) data pertinent to SIC were acquired from the Gene Expression Omnibus (GEO) database. After detailed retrieval, two datasets, GSE229925 and GSE190856, were selected for further analysis. Both studies employed murine models as the primary research organisms, with the former providing bulk RNA-seq data of the mouse hearts 24 hours after cecum ligation and puncture (CLP), and the latter offering scRNA-seq data of immune cells in mouse hearts at steady state and on days 3, 7, and 21 after CLP. For the ensuing analyses, Transcripts per kilobase million (TPM) values were extracted and genes displaying an average expression level below 0.1 were filtered out.

### Bulk RNA-seq data process

We employed the limma package for differential gene expression analysis[9]. The adjusted p-value (Padj) was calculated using the Benjamini-Hochberg (BH) method to reduce the false positive rate. The criteria for identifying significantly differential expressed genes were that Padj <0.05 and |log2FC| >1 (FC, fold change). Gene Ontology (GO) and Kyoto Encyclopedia of Genes and Genomes (KEGG) pathway enrichment analysis were performed using the R package clusterProfiler[10]. Immune cell abundances were estimated using the R package CIBERSORT[11]. Protein-protein interaction (PPI) networks based on the identified differentially expressed genes (DEGs) were constructed using the STRING database[12], and gene pairs with composite scores >0.7 were imported into the Cytoscape software for further analysis. Hub genes related to SIC were then identified using the cytoHubba plugin in the Cytoscape software[13].

### scRNA-seq data process

The GSE190856 scRNA-seq dataset was analyzed using the R package Seurat[14]. To ensure data quality, we initially performed quality control (QC) by retaining cells with less than 10% mitochondrial gene content and genes expressed in at least three cells within an expression range of 200 to 7000. We then identified a set of highly variable genes for further study, with the number of such genes set at 2000. To address batch effects presented in the data from the different samples, we employed the “Harmony” package. Cell clusters were then generated using the “FindClusters” and “FindNeighbors” functions, and “t-SNE” method was used to visualize these clusters. Cell annotation was conducted based on the marker genes associated with different cell types, including macrophages (*Adgre1, Fcgrt, Timd4, Retnla, Lyve1, Cd163, Folr2, Ccr2, H2-Aa,* and *H 2 - E*)*b*, *1* monocytes (*Plac8, Fn1, Ace, Itgal, Napsa, Gngt2,* and *Chil3*), neutrophils (*S100a8/a9, Retnlg, Ifitm1, Lcn2, Ngp,* and *Hp*), natural killer (NK)/T cells (*Cd3e, Klrk1, Ccl5, Gzma, Il1rl1, Gata3,* and *Il7r*), B cells (*Igkc, Ly6d, Ebf1, Cd79b, Ms4a1,* and *Cd79a*) and cycling cells (*Mki67, Stmn1, Top2a, Ube2c,* and *S t m n*). *1*The macrophages were further subdivided into three subsets. The crosstalk among various immune cells were analyzed and then visualized using the R package CellChat[15].

### Cell culture and treatment

The mouse monocyte macrophage leukemia cell line (RAW 264.7) (purchased from the Chinese Academy of Sciences Shanghai Cell Bank, Shanghai, China) was cultivated in DMEM medium (Gibco, USA) supplemented with 10% fetal bovine serum (FBS, Gibco, USA), 100 U/mL penicillin, and 100 μg/mL streptomycin (Solarbio, China). The mouse bone marrow-derived macrophages (BMDMs) were extracted from 5-week-old male C57BL/6J mice and cultivated in RPIM 1640 medium (Gibco, USA) containing 10% FBS (Gibco, USA), 100 U/mL penicillin, 100 μg/mL streptomycin (Solarbio, China), and 50 ng/mL Macrophage-Colony Stimulating Factor (M-CSF, Novoprotein, China). The mouse cardiac microvascular endothelial cells (MCMECs) (purchased from Procell Life Science & Technology Co., Ltd., Wuhan, China) were cultivated in the complete medium for MCMECs (Procell Life Science & Technology Co., Ltd., China). The above cells were cultured in a 37°C atmosphere with 5% CO2.

### SIC murine model and treatments

All animal studies were conducted in accordance with ethical standards and were approved by the Institutional Animal Care and Use Committee of Jiangsu Province Hospital. 8-week-old male C57BL/6J mice were purchased from Jiangsu Huachuang Xinnuo Pharmaceutical Technology Co., Ltd, Taizhou, China and housed in Specific Pathogen Free (SPF) facilities with 23 ± 1 °C temperature and 12/12 hours light-dark cycle. LPS (Cat# L2630, Sigma, USA) was injected intraperitoneally at a dose of 15 mg/kg to simulate sepsis. CD11b (ITGAM) neutralizing antibody (Cat# BE0007, Bio X Cell, USA) was injected intraperitoneally at a dose of 100 μg per mouse for 2 consecutive days and 1 hour prior to LPS modelling. An equal volume of saline was injected into the sham group. Five mice from the LPS group and five from the LPS + neutralizing antibody group underwent echocardiographic examination 24 hours after modeling, followed by euthanasia to collect heart tissues for subsequent assays. Additionally, ten mice from the LPS group and ten from the LPS + neutralizing antibody group were continuously monitored up to day 10 to observe the mortality of the model mice throughout the entire course of SIC.

### Echocardiography

After isoflurane inhalation anesthesia, transthoracic echocardiography was carried out on mice using the Vevo® 3100 high-resolution microsound equipment (Fujifilm Visual Sonics, Canada). An experienced animal experimenter who was blind to the group assignments performed the ultrasound scan. The left ventricular long axis section’s M-mode pictures were used to record the left ventricular ejection fraction (EF) and fraction of short-axis shortening (FS).

### RNA extraction and RT-qPCR

The cardiac tissues, macrophages, and MCMECs were subjected to RNA extraction using the TRIzol Reagent (Cat# 15596026, Invitrogen, USA). The HiScript®R II Q RT SuperMix for qPCR (Cat# R222-001, Vazyme, China) was used to synthesize the cDNA, and the RT-qPCR was performed using ChamQ SYBR qRCR Master Mix (High ROX Premixed) (Cat# Q341-02-AA, Vazyme, China) by the LightCycler® 480 System (Roche, Switzerland) according to the manufacturer’s instructions. Then, the relative expression of genes was calculated by the 2-ΔΔCT method. The primer sequences are displayed as follows: mouse *Itgam* Forword (CTTTGGGAACCTCCGACCAG), mouse *Itgam* Reverse (CACCAAAGTGTCCAAGCCCA), mouse *Bnp* Forword (GAAGGACCAAGGCCTCACAA), mouse *Bnp* Reverse (ACTTCAGTGCGTTACAGCCC), mouse *Icam-1* Forword (GTGATGCTCAGGTATCCATCCA), mouse *Icam-1* Reverse (CACAGTTCTCAAAGCACAGCG), mouse *II-1* Forword (TGCCACCTTTTGACAGTGATG), mouse *Il-1β* Reverse (TGATGTGCTGCTGCGAGATT), mouse *II-6* Forword (GACAAAGCCAGAGTCCTTCAGA), mouse *Il-6* Reverse (TGTGACTCCAGCTTATCTCTTGG), mouse *Mcpt* Forword (GAGGACAGATGTGGTGGGTT), mouse *Mcpt-1* Reverse (AGGAGTCAACTCAGCTTTCTCTT).

### Statistical Analysis

Statistical analyses were conducted using the GraphPad Prism V 8.3.0 software. Normality of the data was assessed using the Shapiro-Wilk test. For data adhering to a normal distribution, comparisons between two groups were performed using an unpaired t-test with Welch’s correction. In cases of non-normal distribution, a standard unpaired t-test was utilized. For the analysis of survival data, Kaplan-Meier survival curves were generated, with intergroup differences evaluated using the log-rank (Mantel-Cox) test. All statistical tests were two-tailed, with a p-value threshold of less than 0.05 set for establishing statistical significance.

## Results

### Identification of the hub genes related to SIC in the bulk RNA seq dataset

Through the analysis of the GSE229925 dataset, we identified 149 significantly up-regulated genes and 43 significantly down-regulated genes in CLP group when compared with the sham group (Figure 1A). Through enrichment analysis (GO and KEGG), we found that in SIC terms related to activation of immune cells (e.g. leukocyte migration, cell activation involved in immune response) and releasion of chemokines (e.g. TNF signaling pathway, IL−17 signaling pathway) were prominently enriched, suggesting a higher inflammatory state in the cardiac tissue of SIC (Figures 1B-1D). We then conducted CIBERSORT analysis to unveil the immune cell composition within the cardiac tissue of SIC mice. The result indicated a significantly higher infiltration of monocytes in the CLP group than in the sham group (Figure 1E). A PPI network was constructed using the STRING database and then imported into the Cytoscape software, fifteen hub genes including Vav1, Cxcl1, Itgam, Selp, Icam1, Ccl2, Cd44, Stat3, Spi1, Itgb2, Hp, Fos, Timp1, Lcn2, and Sele were identified using cytoHubba.

**Figure 1:**
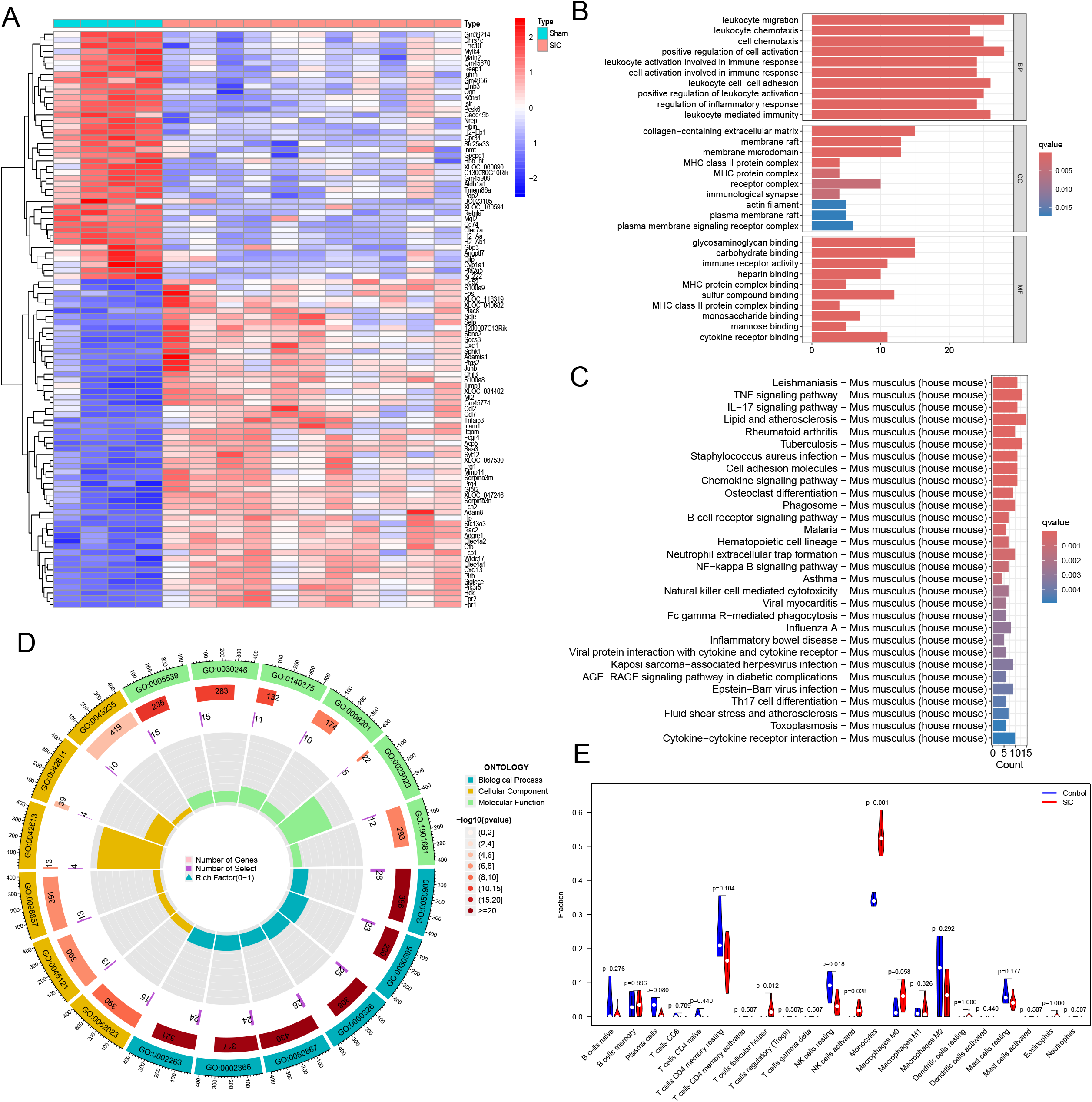
Comprehensive gene expression analysis using the GSE229925 dataset. A. Differential gene expression in the CLP group compared to the sham group, highlighting 149 up-regulated and 43 down-regulated genes. B-D. Enrichment analysis (GO and KEGG) illustrating the predominant activation of immune cells and chemokine pathways, such as TNF and IL-17 signaling. These enriched terms are indicative of an elevated inflammatory state in the cardiac tissue of SIC mice. E. CIBERSORT analysis showing the comparative immune cell composition in the cardiac tissue of SIC mice. Notably, a higher infiltration of monocytes is evident in the CLP group as opposed to the sham group.

### The expression of hub genes in different cardiac immune cells during SIC

We further explore the expression profiles of the fifteen hub genes in different cardiac immune cells using the scRNA-seq data (GSE190856). Single-cell transcriptomic data from mouse hearts at steady state and on days 3, 7, and 21 after CLP were separately analyzed. Immune cell clustering was executed, followed by their categorization into six primary clusters, namely macrophages, monocytes, neutrophils, natural killer (NK)/T cells, B cells and cycling cells. Macrophages emerged as the most prevalent cell type. Upon comparing the alterations in immune cell composition at each time point, we observed a decrease in macrophage counts on day 3 after CLP, followed by an increase on days 7 and 21. Monocytes infiltration was not significant at steady-state. However, in alignment with the results from bulk RNA sequencing, we detected monocyte infiltration on day 3 post-CLP, which peaked on day 7 and gradually diminished by day 21. Additionally, a marked increase in neutrophil infiltration was observed on day 3 after CLP (Figure 2A). Among the fifteen hub genes, *Igtam*, widely acknowledged as a macrophage marker, demonstrated a notable upsurge in expression within macrophages after CLP (Figure 2B). Given that cardiac macrophages can be classified into various subtypes according to their biomarkers expression, functional roles, and embryonic origins, we proceeded to examine the variability in *Igtam* expression across these distinct macrophage subclusters. By utilizing specific molecular markers delineated in prior studies, we categorized cardiac macrophages into three distinct subclusters (MAC1, MAC2, MAC3). MAC1 was characterized by high expression of *Timd4*, *Retnla*, *Lyve1*, *Cd163*, and *Folr2*, representing the self-renewing TLF^+^ resident macrophage subtype in the myocardium[16, 17]. MAC2 corresponded to the MHC-II^hi^ macrophages, they can be partially supplemented by circulating monocytes, but does not undergo continuous renewal[16, 17]. MAC3 represented the CCR2^+^ macrophages, they also expressed high levels of antigen-presentation genes (*H2-Aa* and *H2-Eb1*) and could be rapidly replenished by monocytes [16–18]. Our findings indicated that at steady state, *Igtam* expression was predominantly observed in the MAC1 and MAC3 clusters. Notably, *Igtam* expression was significantly upregulated in the MAC2 cluster after CLP, suggesting that the increased *Igtam* expression on MAC2 might be associated with the progression of SIC (Figure 2C).

**Figure 2:**
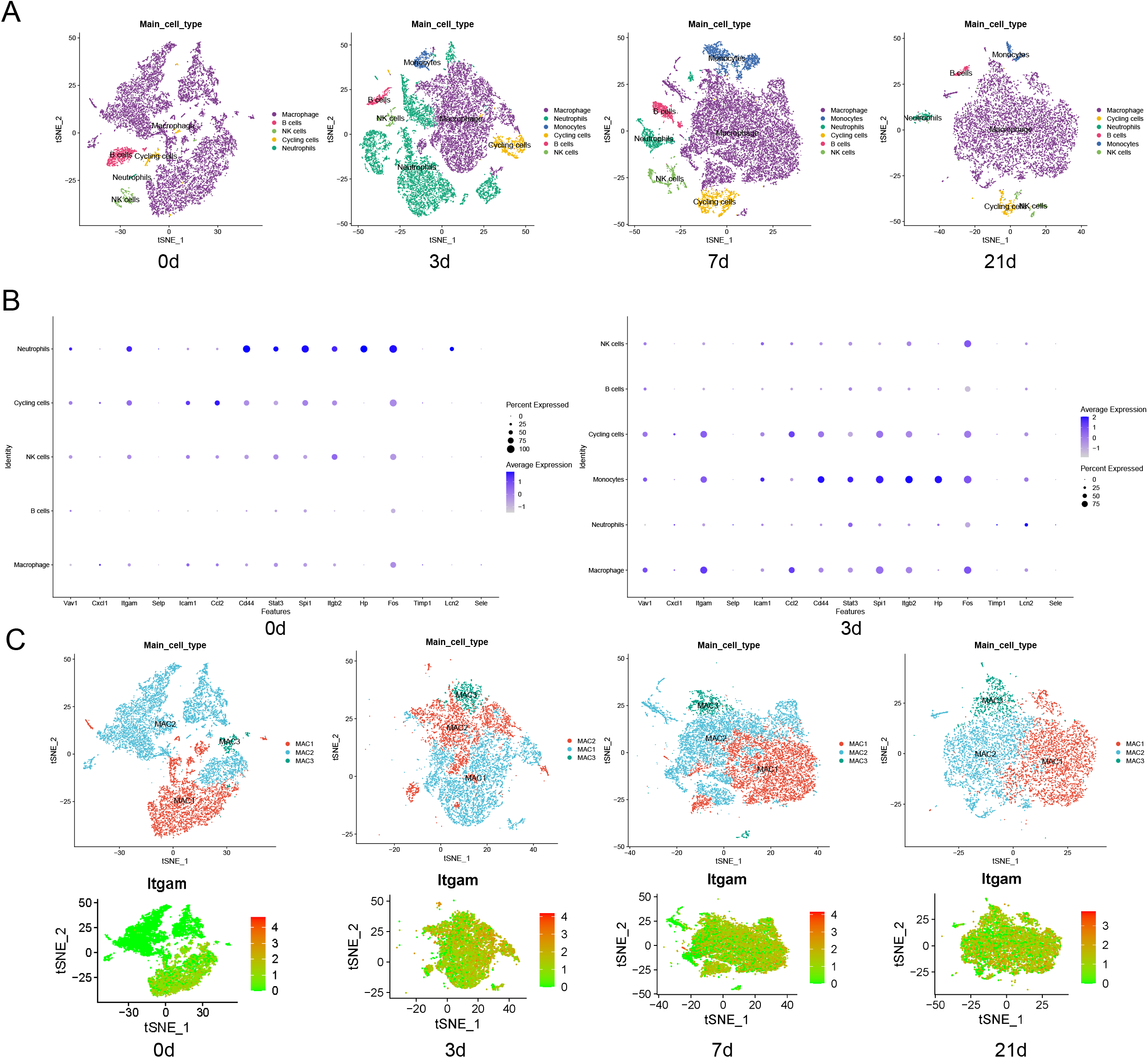
Dynamics of immune cell populations and expression of hub genes in SIC. A. The t-SNE plots show the distribution and dynamics of various immune cell types (macrophages, B cells, neutrophils, NK cells, monocytes, and cycling cells) in the heart tissue at different time points (0, 3, 7, and 21 days) post-cecum ligation and puncture (CLP). B. A detailed view of the macrophage subsets (MAC1, MAC2, MAC3) at each time point, highlighting their relative proportions and transitions during the course of SIC. C. The expression levels of identified hub genes (including Vav1, Cxcl1, Itgam, Selp, Icam1, Ccl2, Cd44, Stat3, Spi1, Itgb2, Hp, Fos, Timp1, Lcn2, and Sele) within the different macrophage subsets, providing insights into the molecular mechanisms driving their roles in the pathogenesis of SIC.

### The role of *Itgam* in immune cell communication during SIC

As ITGAM typically combines with the integrin subunit beta 2 (ITGB2) to form a leukocyte-specific integrin also known as macrophage receptor 1, we further investigated the role of *Itgam* in facilitating communication among diverse immune cells by using the “Secreted Signaling” and “Cell-Cell contact” databases curated in CellChat. Employing the “Secreted Signaling” database revealed a significant activation of the C3 − (Itgam+Itgb2) ligand-receptor pair after CLP, contributing to the signaling from monocytes to macrophages, monocytes to monocytes, and monocytes to cycling cells (Figure 3A). On the other hand, when using the “Cell-Cell contact” database, we found that the ICAM signaling, which was known for regulating leukocyte recruitment from circulation to sites of inflammation, was upregulated through the Icam1 − (Itgam+Itgb2) ligand-receptor pair. Notably, it was observed that monocytes, a subset of immune cells not typically residing in the heart at steady state, exhibited a robust response to macrophages post-CLP, mediated by the ICAM signaling. Additionally, our findings demonstrated that primarily four types of immune cells, namely monocytes, macrophages, NK cells, and cycling cells, engage in cell-cell contact communication through this ligand-receptor pair, as illustrated in Figure 3B.

**Figure 3:**
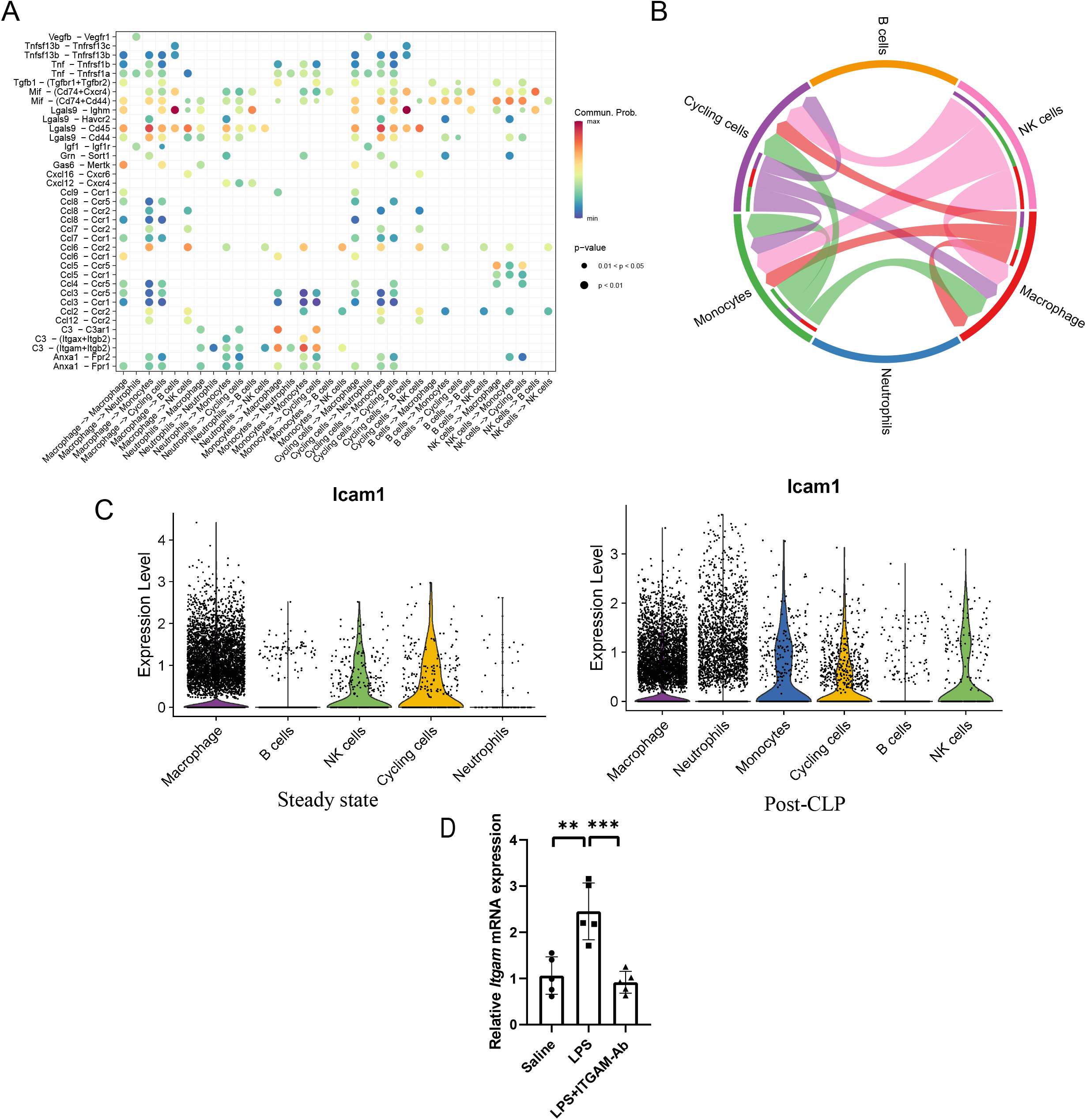
Elucidating the role of ITGAM in immune cell communication and expression during SIC. A. Activation of the C3 − (Itgam+Itgb2) ligand-receptor pair post-CLP, highlighting enhanced signaling pathways among monocytes, macrophages, and cycling cells. This diagram illustrates the pivotal role of ITGAM in modulating intercellular communication in the context of SIC. B. Representation of cell-cell contact communication facilitated by the ICAM1 − (Itgam+Itgb2) ligand-receptor pair, predominantly involving monocytes, macrophages, NK cells, and cycling cells. This figure underscores the significant upregulation of ICAM signaling post-CLP. C. Analysis of *Icam1* expression across various immune cell types at steady state and following CLP. D. Graphical representation of the pronounced elevation in *Icam1* expression within myocardial tissues post-CLP, as derived from the GSE229925 dataset.

Analogous to *Itgam*, *Icam1* was also identified as one of the 15 hub genes associated with SIC within this study. We further examined the expression of *Icam1* in various immune cells following CLP but did not observe a significant upregulation of its expression akin to that of *Itgam* (Figure 3C). To comprehensively evaluate the global expression profile of the *Icam1* gene in cardiac tissues during SIC, the GSE229925 dataset was utilized and the analytical outcomes revealed a pronounced elevation in *Icam1* expression within the myocardial tissues post-CLP (Figure 3D). Given that *Icam1* is typically expressed in immune cells and vascular endothelial cells, it is posited that an augmentation in *Icam-1* expression occurs within endothelial cells during SIC.

### Upregulation of *Itgam* in macrophages and *Icam-1* in endothelial cells following LPS stimulation

We utilized LPS to stimulate the RAW 264.7 mouse monocyte-macrophage leukemia cell line, as well as primary mouse bone marrow-derived macrophages (BMDM), aiming to replicate the septic state in an in vitro setting. Subsequent qPCR revealed a notable elevation in *Itgam* expression in both RAW 264.7 cells and primary macrophages following LPS stimulation (Figure 4A). To further verify the hypothesis of *I c a m – 1* upregulation in endothelial cells during SIC, mouse cardiac microvascular endothelial cells (MCMEC) were subjected to LPS stimulation, resulting in a marked increase in *Icam-1* expression (Figure 4B).

**Figure 4:**
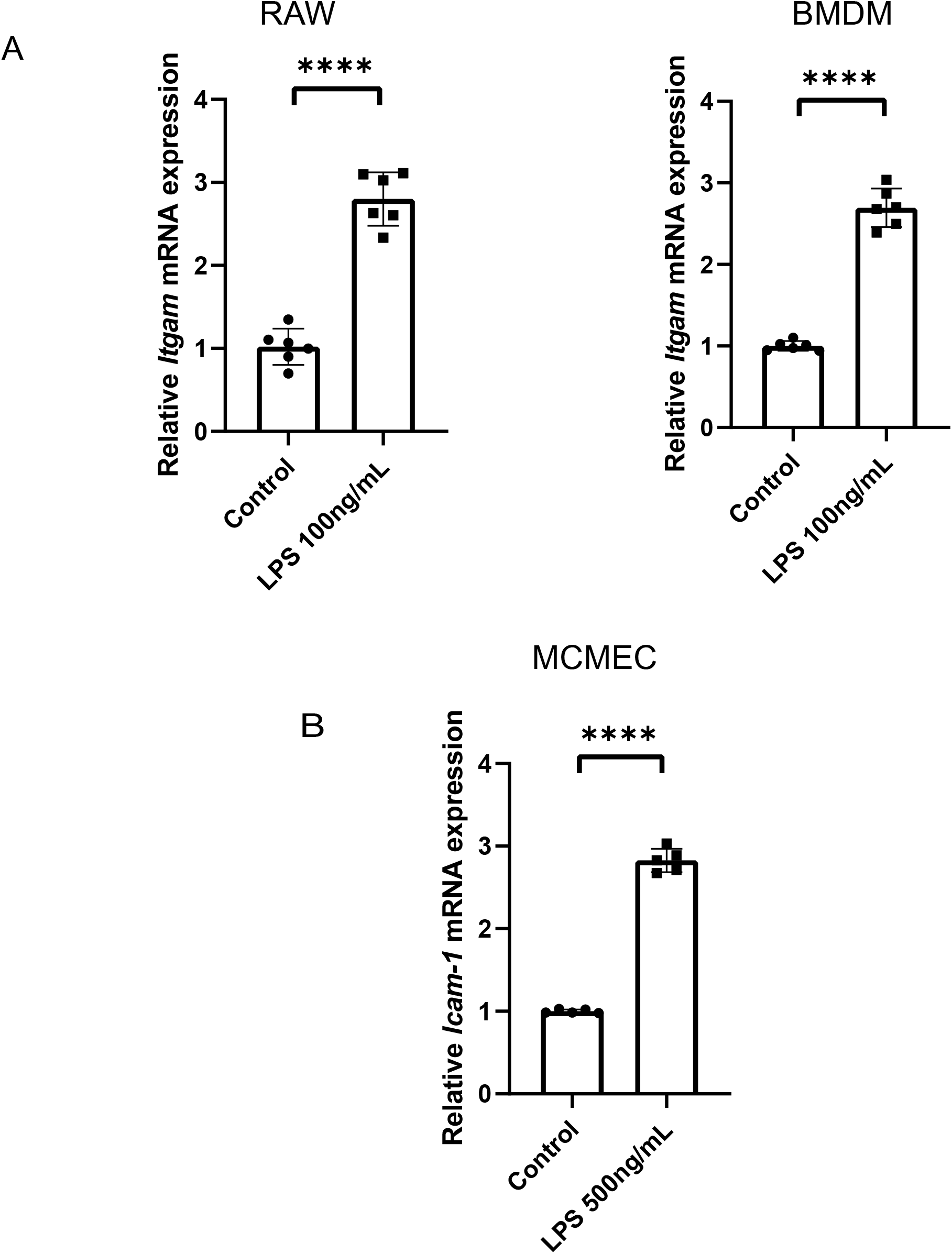
Upregulation of *Itgam* in Macrophages and *Icam-1* in Endothelial Cells Following LPS Stimulation. A. The relative *Itgam* mRNA expression in RAW 264.7 cells and primary mouse bone marrow-derived macrophages (BMDM) post-LPS stimulation. The data illustrates a significant increase in *Itgam* expression in both cell types. B. The relative *Icam-1*mRNA expression in mouse cardiac microvascular endothelial cells (MCMEC) after LPS stimulation. The marked elevation in Icam-1 expression upon exposure to LPS (500ng/mL) underscores the responsiveness of endothelial cells in the context of SIC.

### Impact of ITGAM neutralization on macrophages recruitment and cardiac outcomes during SIC

To further elucidate the role of ITGAM in SIC, we administered an ITGAM-specific neutralizing antibody to mice to inhibit ITGAM function in vivo before giving the LPS intraperitoneal injection for modelling. The results demonstrated a diminution in *Itgam* expression within myocardial tissues following the administration of the neutralizing antibody (Figure 5A). This observation, coupled with our cellular experiments showing increased *Itgam* expression in macrophages and elevated *I c a m* e*-*x*1*pression in endothelial cells following LPS stimulation, indicated a marked reduction in the infiltration of ITGAM-positive monocytes/macrophages within myocardial tissues after neutralizing antibody administration. Concurrently, the expression of *Bnp* mRNA, a marker of heart failure, was lower in the neutralizing antibody group compared to the sham group 24 hours after LPS modeling. And there was also an opposite trend observed in the expression of some inflammation-related genes, such as *Il-6*, *Il-1β*, and *Mcpt-1* (Figure 5B). Similarly, results from small animal echocardiography echoed these findings, suggesting that inhibiting ITGAM exerts a cardioprotective effect in the early stage of SIC (Figure 5C). However, although the mice in the neutralizing antibody group exhibited better cardiac function in the early stages of SIC, the overall mortality rate was higher in the neutralizing antibody group throughout the course of the disease, with most of these deaths occurring in the mid to late stages of SIC (Figure 5D).

**Figure 5:**
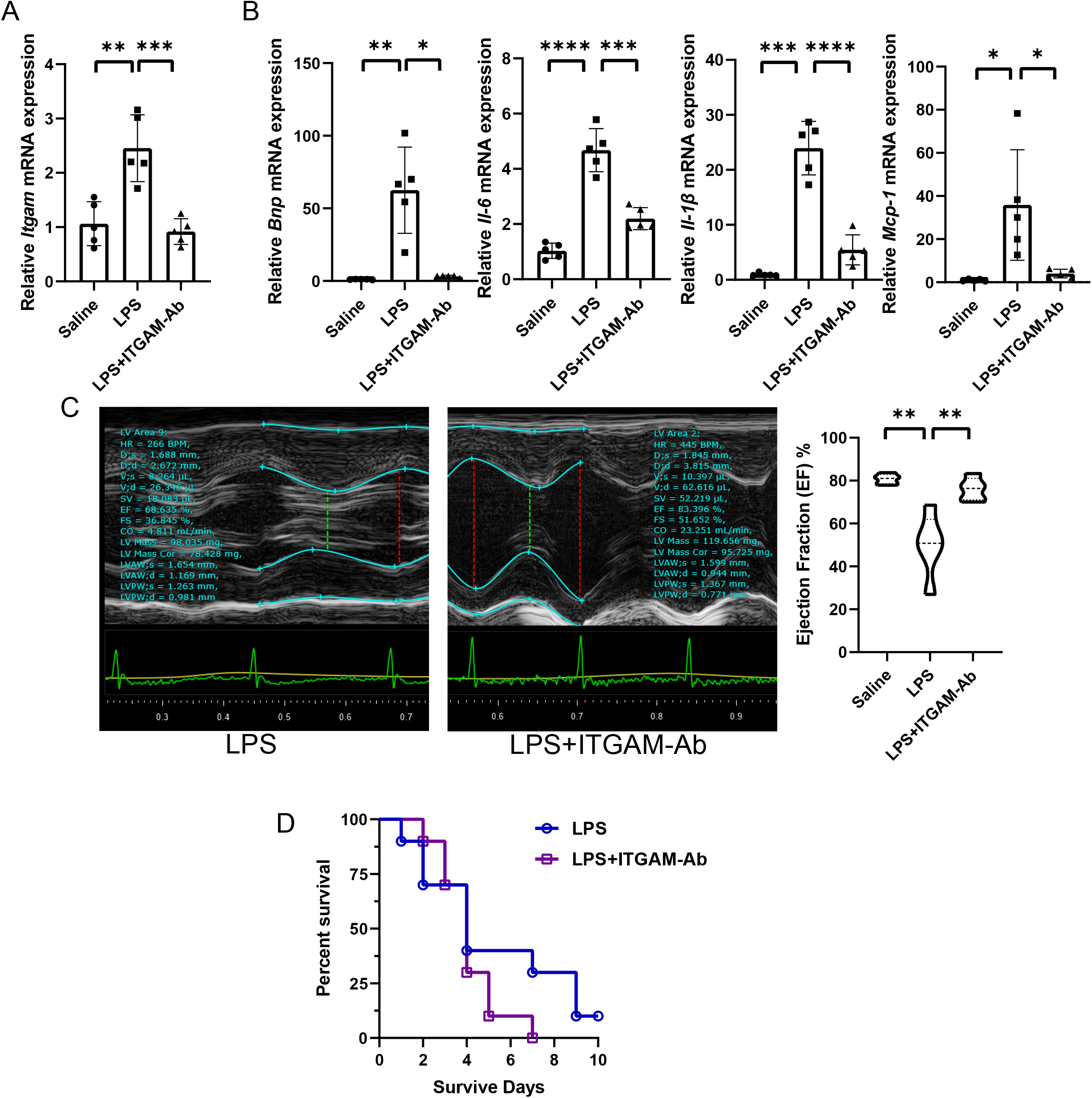
The impact of ITGAM neutralization on macrophage recruitment and cardiac function during SIC. A. The relative expression of *I t g a* m*m*RNA in myocardial tissues following the administration of ITGAM-specific neutralizing antibody. B. The relative expression levels of *Bnp* mRNA and other inflammation-related genes (Il-6, Il-1β, and Mcpt-1) in the myocardial tissues. The neutralizing antibody group exhibits lower expression of *Bnp* mRNA and higher expression of inflammatory markers, indicating reduced cardiac injury in this group during the early stage of SIC. C: Illustrates the results from small animal echocardiography, particularly the ejection fraction percentage, indicating a cardioprotective effect in the early stage of SIC in the ITGAM neutralizing antibody group compared to the LPS and saline groups. D: The Kaplan-Meier survival curves, showing the percent survival over 10 days. Notably, despite improved early cardiac function, the overall mortality rate was higher in the ITGAM neutralizing antibody group, particularly in the mid to late stages of SIC.

## Discussion

In the present research, comprehensive bioinformatics analysis afforded profound insights into the dynamic alterations in gene expression and immune cell composition within cardiac tissue during SIC. Irrespective of being in a steady state or subsequent to CLP modeling, macrophages consistently represented the predominant immune cell constituent within the cardiac milieu, accompanied by a pronounced upregulation of *Itgam* post-CLP, delineating their potential contributory role in the progression of SIC. Additionally, our investigation into immune cell communication underscored the critical role of ITGAM and ICAM-1 in orchestrating interactions among monocytes, macrophages, NK cells, and cycling cells during SIC. These findings not only deepen our understanding of the molecular mechanisms underpinning SIC but also potentially open avenues for targeted therapeutic interventions.

Resident cardiac macrophages (RCMs) have been shown to play a protective role in maintaining cardiac structure and function in conditions like myocardial infarction, heart failure, and myocarditis[16]. In the context of SIC, the researchers of GSE190856 revealed that TREM2^hi^ resident macrophages scavenge cardiomyocyte-ejected dysfunctional mitochondria, thus preventing excessive inflammation and sustaining normal cardiac function[18]. Drawing on prior studies, we categorizes macrophages into three distinct subtypes in our analysis of the GSE190856 dataset: the MAC1 was TLF^+^ macrophages (*Timd4*, *Lyve1*, *Folr2*), they were evolutionary conserved and self-renewing, mainly involved in cellular transport and endocytosis, with TREM2 highly expressed in this particular subset; the MAC2 was MHC-II^hi^ macrophages, they were partially replaced by circulating monocytes (∼25%), while previous studies have shown high TREM2 expression in this subset in lung, liver, and kidney[17], such a trend was not significant in heart in the GSE190856 dataset[18]; the MAC3 was CCR2+ macrophages, they were closely related to circulating monocytes and involved in immune processes such as Nlrp3 inflammasome activation and Il1b production. Serving as a key macrophage surface marker, *Itgam* was highly expressed in MAC1 and MAC3 at steady state, yet notably absent in MAC2. However, during the course of SIC, a significant upsurge in *Itgam* expression within the MAC2 subset was observed, pointing to its dynamic role in the disease’s progression.

It has been observed that the monocytes infiltrating to the heart gradually adopt the transcriptional profiles synonymous with each distinct RCM subset over time[17]. To determine whether the augmented *Itgam* expression in MAC2 stemmed from the transitioning of recruited circulating monocytes/macrophages into MAC2 or a surge in *Itgam* expression within the inherent resident MAC2, we first demonstrated that peripheral macrophages (using the RAW cell line and primary mouse bone marrow cells) exhibited increased *Itgam* expression following LPS stimulation. Concurrently, endothelial cells exhibited a marked elevation in *Icam-1* expression under LPS stimulation. Following this, in a mouse model of SIC, qPCR analysis disclosed a notable upregulation of *Itgam* expression in the myocardium. However, administering an ITGAM specific blocking antibody during the modeling process effectively neutralized this trend. These findings imply that while MAC2 predominantly self-replenishes at steady state as per prior studies[17], during SIC however, peripheral monocytes/macrophages are likely significantly recruited to the heart via the upregulated expression of ICAM-1 on endothelial cells. These monocytes/macrophages then differentiated to MAC2, and thus amplifying the cohort of MAC2 with high *Itgam* expression. Coupled with cell communication analysis indicating intensified interactions between macrophages and other immune cells through the ICAM1 − (ITGAM+ITGB2) ligand-receptor pair during SIC, we deduce that MAC2 exerts a crucial role in the pathogenesis of SIC via *Itgam*.

ITGAM, also known as CD11b, synergizes with the beta 2 chain (ITGB2) to constitute a leukocyte-specific integrin, commonly recognized as macrophage receptor 1, complement receptor 3, or Mac-1. This integrin is documented as essential for facilitating the adhesion of neutrophils and monocytes to activated endothelium through ICAM-1 and plays a pivotal role in the phagocytosis of complement-coated particles[19–22]. This could elucidate the observations noted at the onset stage of SIC: circulating monocytes/macrophages, after being stimulated by inflammatory stimulation, exhibited an upsurge in *Itgam* expression. These cells were actively recruited to the heart through ICAM-1, which was highly expressed on cardiac endothelial cells. Subsequently, their interactions with various immune cells were amplified through the C3 − (ITGAM+ITGB2) and the ICAM1 − (ITGAM+ITGB2) ligand-receptor pairs, exacerbating the inflammatory response within the heart and consequently impacting cardiac functionality. Meanwhile, these circulating macrophages also transformed into the MAC2 phenotype during this process. Although the precise functions of MHC-II^hi^ macrophages beyond antigen presentation remain to be fully elucidated, multiple pieces of evidence point to their immunoregulatory and cardioprotective roles in situations like cardiac pressure overload and myocardial infarction [23, 24]. Therefore, in the middle and later stages of SIC, these cells might be key factors in promoting disease alleviation. In fact, the function of Mac-1 is also perennially perceived as possessing a dual nature. In a research on thrombotic glomerulonephritis, Mac-1 was pinpointed as a critical molecular nexus interlinking inflammation and thrombosis, influencing neutrophil recruitment, endothelial injury, and the development of thrombosis, while also playing a role in maintaining vessel wall integrity against inflammation-induced damage[25]. In autoimmune diseases, besides its role in promoting inflammation through leukocyte recruitment and leukocyte cytotoxic functions, Mac-1 can also exert anti-inflammatory effects through mechanisms such as apoptotic cell clearance, inhibition of B cell activation, induction of tolerogenic responses by dendritic cells, and T-cell inhibition[21]. In this study, we found that during SIC, the expression of *Itgam* in MHC-II^hi^ macrophages within the heart is significantly increased. In light of the dual role of Mac-1, we postulate that in the early stages of SIC, circulating monocytes/macrophages are recruited to the heart, leading to an excessive inflammatory response that affects cardiac function. However, as these cells undergo transformation into MHC-II^hi^ macrophages upon cardiac entry, they are posited to exert immunoregulatory and anti-inflammatory effects through Mac-1 and other related signaling pathways, thereby improving cardiac function during the mid to late stages of SIC.

Despite the comprehensive nature of our findings in elucidating the mechanisms and cellular dynamics of SIC, this study is not without its limitations. Firstly, our reliance on bioinformatics analysis and specific datasets, while providing a robust foundation for our hypotheses, also means that our conclusions are inherently tied to the limitations of these datasets. The extrapolation of these findings to broader contexts should be approached with caution. Furthermore, while our study offers valuable insights into the role of ITGAM and ICAM-1 in macrophages recruitment and cardiac outcomes during SIC, it primarily focuses on the early stages of the disease. The higher mortality observed in the neutralizing antibody group in the later stages of SIC suggests complex underlying mechanisms that our study did not fully explore. This indicates a need for more extensive experimental studies to understand the long-term effects of ITGAM inhibition and the intricate balance between protective and detrimental immune responses in SIC. An additional limitation of our study is the exclusive use of murine models. While mouse models are invaluable in biomedical research for their genetic similarity to humans and their utility in experimental manipulation, there are intrinsic differences between murine and human physiological and immunological systems. Future research endeavors should aim to incorporate human-derived data.

## Conclusion

This study sheds light on the complex role of ITGAM in the progression of SIC, particularly highlighting the reversible nature of myocardial dysfunction. Our findings suggest that during the early stages of SIC, an influx of circulating monocytes/macrophages, exacerbated by ITGAM, intensifies cardiac inflammation, impairing cardiac function. However, as the disease progresses, these cells differentiate into MHC-II^hi^ macrophages (MAC2), potentially reversing the detrimental effects and aiding in the restoration of cardiac function. The dual role of Mac-1 (ITGAM+ITGB2) may also contribute to both the exacerbation and amelioration of cardiac dysfunction. Our research offers vital insights into the molecular mechanisms of SIC and underscores the need for further studies to extend these findings from murine models to human clinical scenarios, aiming for more effective therapeutic strategies in sepsis management.

### Declaration of generative AI and AI-assisted technologies in the writing process

During the preparation of this work, the author(s) used ChatGPT Classic to optimize and correct the language expression. After using this tool, the author(s) reviewed and edited the content as needed and take full responsibility for the content of the publication.

## References

[1] K.E. Rudd, S.C. Johnson, K.M. Agesa, K.A. Shackelford, D. Tsoi, D.R. Kievlan, D.V. Colombara, K.S. Ikuta, N. Kissoon, S. Finfer, C. Fleischmann-Struzek, F.R. Machado, K.K. Reinhart, K. Rowan, C.W. Seymour, R.S. Watson, T.E. West, F. Marinho, S.I. Hay, R. Lozano, A.D. Lopez, D.C. Angus, C.J.L. Murray, M. Naghavi, Global, regional, and national sepsis incidence and mortality, 1990-2017: analysis for the Global Burden of Disease Study, Lancet (London, England), 395 (2020) 200–211.

[2] S.M. Hollenberg, M. Singer, Pathophysiology of sepsis-induced cardiomyopathy, Nature reviews. Cardiology, 18 (2021) 424–434.

[3] R.J. Levy, D.A. Piel, P.D. Acton, R. Zhou, V.A. Ferrari, J.S. Karp, C.S. Deutschman, Evidence of myocardial hibernation in the septic heart, Critical care medicine, 33 (2005) 2752–2756.

[4] S.M. Hollenberg, Think locally: evaluation of the microcirculation in sepsis, Intensive care medicine, 36 (2010) 1807–1809.

[5] A. Conway-Morris, J. Wilson, M. Shankar-Hari, Immune Activation in Sepsis, Critical care clinics, 34 (2018) 29–42.

[6] Y. Kakihana, T. Ito, M. Nakahara, K. Yamaguchi, T. Yasuda, Sepsis-induced myocardial dysfunction: pathophysiology and management, Journal of intensive care, 4 (2016) 22.

[7] G. Stanzani, M.R. Duchen, M. Singer, The role of mitochondria in sepsis-induced cardiomyopathy, Biochimica et biophysica acta. Molecular basis of disease, 1865 (2019) 759–773.

[8] D. Brealey, M. Brand, I. Hargreaves, S. Heales, J. Land, R. Smolenski, N.A. Davies, C.E. Cooper, M. Singer, Association between mitochondrial dysfunction and severity and outcome of septic shock, Lancet (London, England), 360 (2002) 219–223.

[9] M.E. Ritchie, B. Phipson, D. Wu, Y. Hu, C.W. Law, W. Shi, G.K. Smyth, limma powers differential expression analyses for RNA-sequencing and microarray studies, Nucleic acids research, 43 (2015) e47.

[10] T. Wu, E. Hu, S. Xu, M. Chen, P. Guo, Z. Dai, T. Feng, L. Zhou, W. Tang, L. Zhan, X. Fu, S. Liu, X. Bo, G. Yu, clusterProfiler 4.0: A universal enrichment tool for interpreting omics data, Innovation (Cambridge (Mass.)), 2 (2021) 100141.

[11] A.M. Newman, C.L. Liu, M.R. Green, A.J. Gentles, W. Feng, Y. Xu, C.D. Hoang, M. Diehn, A.A. Alizadeh, Robust enumeration of cell subsets from tissue expression profiles, Nature methods, 12 (2015) 453–457.

[12] D. Szklarczyk, R. Kirsch, M. Koutrouli, K. Nastou, F. Mehryary, R. Hachilif, A.L. Gable, T. Fang, N.T. Doncheva, S. Pyysalo, P. Bork, L.J. Jensen, C. von Mering, The STRING database in 2023: protein-protein association networks and functional enrichment analyses for any sequenced genome of interest, Nucleic acids research, 51 (2023) D638–d646.

[13] C.H. Chin, S.H. Chen, H.H. Wu, C.W. Ho, M.T. Ko, C.Y. Lin, cytoHubba: identifying hub objects and sub-networks from complex interactome, BMC systems biology, 8 Suppl 4 (2014) S11.

[14] A. Butler, P. Hoffman, P. Smibert, E. Papalexi, R. Satija, Integrating single-cell transcriptomic data across different conditions, technologies, and species, Nature biotechnology, 36 (2018) 411–420.

[15] S. Jin, C.F. Guerrero-Juarez, L. Zhang, I. Chang, R. Ramos, C.H. Kuan, P. Myung, M.V. Plikus, Q. Nie, Inference and analysis of cell-cell communication using CellChat, Nature communications, 12 (2021) 1088.

[16] R. Zaman, S. Epelman, Resident cardiac macrophages: Heterogeneity and function in health and disease, Immunity, 55 (2022) 1549–1563.

[17] S.A. Dick, A. Wong, H. Hamidzada, S. Nejat, R. Nechanitzky, S. Vohra, B. Mueller, R. Zaman, C. Kantores, L. Aronoff, A. Momen, D. Nechanitzky, W.Y. Li, P. Ramachandran, S.Q. Crome, B. Becher, M.I. Cybulsky, F. Billia, S. Keshavjee, S. Mital, C.S. Robbins, T.W. Mak, S. Epelman, Three tissue resident macrophage subsets coexist across organs with conserved origins and life cycles, Science immunology, 7 (2022) eabf7777.

[18] K. Zhang, Y. Wang, S. Chen, J. Mao, Y. Jin, H. Ye, Y. Zhang, X. Liu, C. Gong, X. Cheng, X. Huang, A. Hoeft, Q. Chen, X. Li, X. Fang, TREM2(hi) resident macrophages protect the septic heart by maintaining cardiomyocyte homeostasis, Nature metabolism, 5 (2023) 129–146.

[19] M.S. Diamond, D.E. Staunton, S.D. Marlin, T.A. Springer, Binding of the integrin Mac-1 (CD11b/CD18) to the third immunoglobulin-like domain of ICAM-1 (CD54) and its regulation by glycosylation, Cell, 65 (1991) 961–971.

[20] Z. Li, The alphaMbeta2 integrin and its role in neutrophil function, Cell research, 9 (1999) 171–178.

[21] F. Rosetti, T.N. Mayadas, The many faces of Mac-1 in autoimmune disease, Immunological reviews, 269 (2016) 175–193.

[22] A. Torres-Gomez, C. Cabañas, E.M. Lafuente, Phagocytic Integrins: Activation and Signaling, Frontiers in immunology, 11 (2020) 738.

[23] N.R. Wong, J. Mohan, B.J. Kopecky, S. Guo, L. Du, J. Leid, G. Feng, I. Lokshina, O. Dmytrenko, H. Luehmann, G. Bajpai, L. Ewald, L. Bell, N. Patel, A. Bredemeyer, C.J. Weinheimer, J.M. Nigro, A. Kovacs, S. Morimoto, P.O. Bayguinov, M.R. Fisher, W.T. Stump, M. Greenberg, J.A.J. Fitzpatrick, S. Epelman, D. Kreisel, R. Sah, Y. Liu, H. Hu, K.J. Lavine, Resident cardiac macrophages mediate adaptive myocardial remodeling, Immunity, 54 (2021) 2072–2088.e2077.

[24] J. Yap, J. Irei, J. Lozano-Gerona, S. Vanapruks, T. Bishop, W.A. Boisvert, Macrophages in cardiac remodelling after myocardial infarction, Nature reviews. Cardiology, 20 (2023) 373–385.

[25] J. Hirahashi, K. Hishikawa, S. Kaname, N. Tsuboi, Y. Wang, D.I. Simon, G. Stavrakis, T. Shimosawa, L. Xiao, Y. Nagahama, K. Suzuki, T. Fujita, T.N. Mayadas, Mac-1 (CD11b/CD18) links inflammation and thrombosis after glomerular injury, Circulation, 120 (2009) 1255–1265.

